# Conversion of somatic sex identity in the testis induces female-specific cellular behaviors in the soma and early oocyte specification in the germline

**DOI:** 10.1101/2024.11.03.621503

**Authors:** Tiffany V. Roach, Sneh Harsh, Rajiv Sainath, Erika A. Bach, Kari F. Lenhart

## Abstract

Establishment and maintenance of cellular sex identity is essential for reproduction. Critically, the sex identity of somatic and germline cells must correspond for sperm or oocytes to be produced, with mismatched identity causing infertility in all organisms from flies to humans. In the *Drosophila* testis, Chronologically inappropriate morphogenesis (Chinmo) is required for maintenance of adult male somatic identity. Loss of *chinmo* leads to progressive feminization of the male soma, including adoption of female-specific cell morphologies, tissue organization and gene expression. However, the degree to which this feminized soma in the male engages female-specific cellular behaviors or influences the associated XY germline is unknown. Using extended live imaging, we have visualized the process of male-to-female somatic sex conversion upon *chinmo* loss of function. We find that feminized soma in the testis engage cell behaviors characteristic of ovarian follicle cells (FCs), including female-specific incomplete cytokinesis, as early as one day of *chinmo* inhibition. In the ovary, FCs collectively migrate around the underlying germ cells to establish the elongated shape of the oocyte. Surprisingly, we find that FC-like soma in *chinmo*-depleted testes also engage this female-specific collective cell migration. Critically, migration of FC-like cells in the testis has the same molecular requirements as FC migration in the ovary. Depletion of the basement membrane protein Perlecan or adhesion protein Ecadherin significantly disrupts rotational migration in both the ovary and *chinmo*-depleted testes. Finally, we find that feminized soma non-autonomously alters sex identity of the associated XY germ cells, inducing expression of a protein required for early oocyte specification. Taken together, our work reveals a dramatic transformation of somatic cell behavior during the process of sex conversion and provides a powerful model to study soma-derived induction of oocyte identity.

## Introduction

Correct sex specification and differentiation is essential for development and maintenance of tissue homeostasis^1–3^. In particular, it is essential for soma and germline sex identity to match for proper gamete production^4,5^. Mismatched sex identity can lead to dramatic changes to cell morphologies and gene expression, ultimately causing sterility^6^. Some disorders of sex determination (DSDs) such as Klinefelter (XXY) or Turner (XO) syndrome are caused by improper numbers of sex chromosomes^7,8^. However, other DSDs which do not involve altered sex chromosome numbers but are likely due to issues in sex maintenance pathways remain poorly understood^9^. The *Drosophila* gonad is an ideal system for investigating mechanisms underlying sex maintenance because it is a well characterized sexually dimorphic tissue^10^. Within the adult fly testis and ovary, there are morphologically distinct somatic lineages that engage sex-specific behaviors to facilitate formation of sperm and oocytes. In a wildtype testis, somatic cyst stem cells (CySCs) give rise to daughters, two of which encapsulate differentiating male germ cells (Fig. 1A) for successful sperm production^11–15^. In a wildtype ovary, follicle stem cells produce follicle cells (FCs), which form a rotating epithelium around female germ cell clusters (Fig. 1D) to properly shape the oocyte^16–18^. Recent work suggests these sex-specific characteristics must be actively maintained over the lifetime of the organism^19,20^.

**Figure 1.**
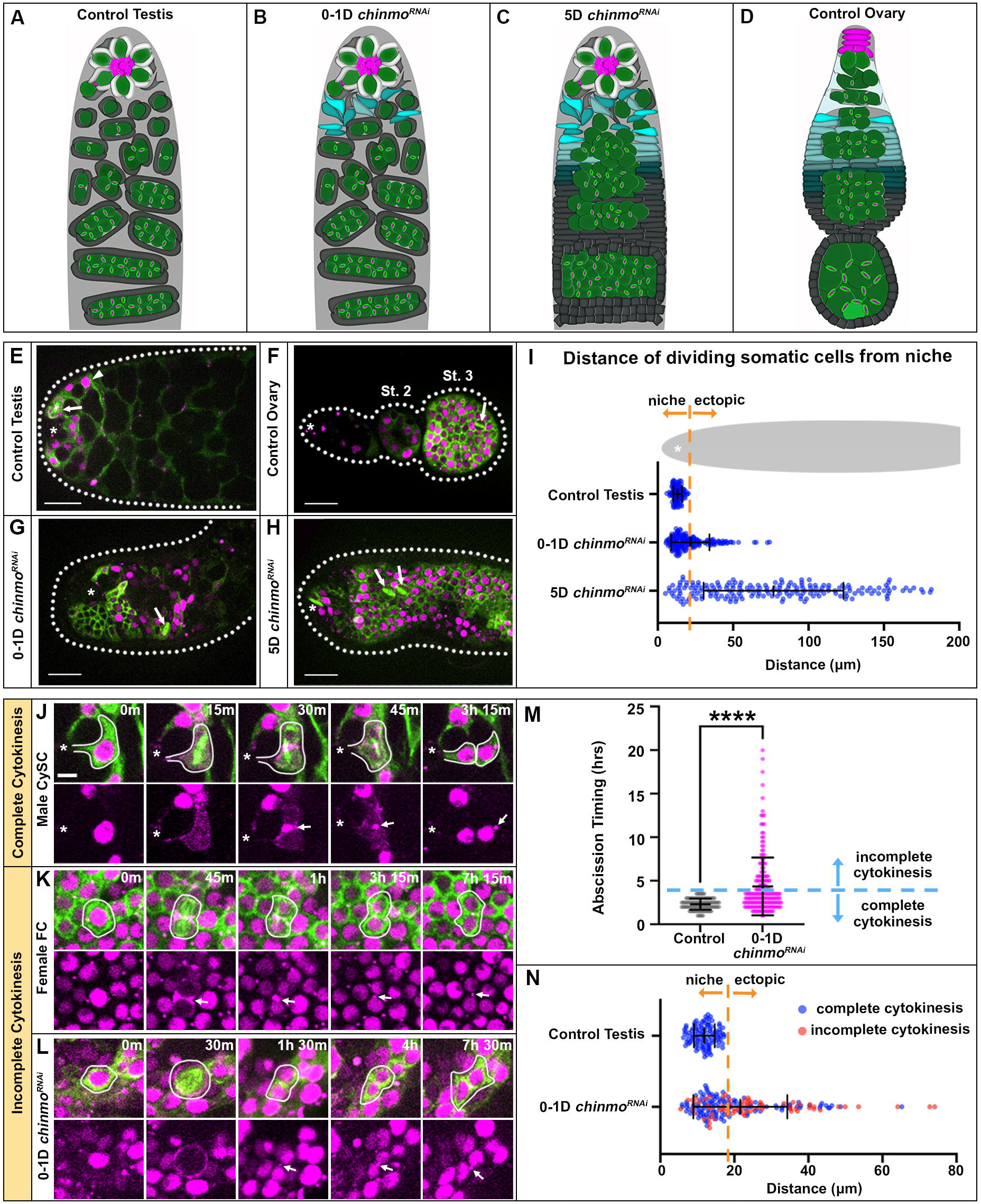
*chinmo*-deficient somatic cells in the testis acquire a female specific cytokinesis program. (A-D) Diagrams showing soma-germline organization in a (A) wildtype testis, (B) 0-1D *chinmo*^*RNAi*^ testis and (C) 5D *chinmo*^*RNAi*^ testis, showing progressive feminization of somatic cell morphology, and a (D) wildtype ovary. (E-H) Stills from time-lapse imaging of somatic cells expressing UAS tubulin::GFP (green) and UAS anillin::RFP (magenta) in the anterior region of a (E) control testis (arrowhead indicates a quiescent cyst cell), (F) control ovary, (G) 0-1D *chinmo*^*RNAi*^ testis, and (H) 5D *chinmo*^*RNAi*^ testis (arrows indicate dividing cells). Scale bar: 20 μm. (I) Quantification of the distance of all dividing somatic cells from the niche (*n* ≥ 113 cells in 10 testes). (J-L) Time-lapse of somatic tubulin (green) and anillin (magenta) during division and cytokinesis in (J) male CySCs, (K) female FCs, and (L) 0-1D *chinmo*^*RNAi*^ soma (arrows point to midbodies). Asterisks indicate niches. White lines outline somatic cells. (M) Quantification of cytokinesis timing through abscission (*n* ≥ 113 cells in 10 testes). \*\*\*\**p* <0.0001 (non-parametric Mann–Whitney *U*-test). (N) Data from panel I shown in false color to indicate somatic cells that completed cytokinesis (blue) or exhibited incomplete cytokinesis (red). All experiments *n* ≥ 2 trials. Scale bar: 5 μm (for J-L). Each image is 1-5 z-slices.

In the *Drosophila* testis, Chronologically inappropriate morphogenesis (Chinmo) is a transcription factor known to maintain male somatic sex identity^19,21,22^. *chinmo*-deficient somatic cells have been shown to “feminize”, turning on female-specific markers and undergoing massive morphological changes reminiscent of female FCs (Fig. 1B-C). However, whether the induction of female gene expression and transformation to FC-like morphologies coincides with acquisition of female specific cellular behaviors has not been addressed. Female somatic cells also non-autonomously regulate female germline identity during development, and proper female sex identity in developing germ cells is required for adult oogenesis (reviewed in^23,24^). However, few molecular details are known about somatic control over the female germline during development^25,26^, and no details are known about whether somatic signals control germline identity in the adult. Studying adult XY germ cells surrounded by a feminized adult soma could yield new insights.

To this end, we have established an extended live-cell imaging technique to visualize somatic and germ cell behaviors within *ex vivo* cultured gonads. Using this model system, we interrogated real time dynamics of sex converted testis soma and discovered progressive acquisition of female-specific cell behaviors. Surprisingly, in fully feminized testes with a complete FC-like epithelium, somatic cells engage rotational migration around the germline. We show that functional rotation depends on expression of adherens and ECM proteins by FCs and FC-like somatic cells. Furthermore, we find evidence that sex-converted testis soma alters the behavior and identity of underlying adult XY germ cells. Finally, our data uncovers a unique model for parsing the requirement of soma versus germline-derived signaling in establishing early oocyte identity in the germline.

## Results and Discussion

### *chinmo*-deficient somatic cells acquire a female-specific cytokinesis program in the testis

In wildtype testes, only CySCs at the niche divide while their differentiating progeny remain quiescent as they encyst germ cells (Fig. 1E,I). In ovaries, follicle stem cells residing outside the niche give rise to mitotically active FCs that form an epithelium around germ cells (Fig. 1F). Previous work has identified ectopic divisions of *chinmo*-deficient somatic cells as an early indication of male-to-female sex conversion^21^. Indeed, our live imaging reveals somatic cell divisions outside of the niche as early as 0-1D of adulthood (0-1D tj>*chinmo* RNAi, hereafter *chinmo*^*RNAi*^) (Fig. 1G, I). At the start of imaging, the organization of soma is identical to wildtype testes but begins undergoing morphological changes over a 24-hour (hr) period. Through direct quantification of cell divisions, we show significant somatic expansion similar to that of pre-FCs, which divide rapidly to form early egg chambers^27^. By 5D post-eclosion (5D *chinmo*^*RNAi*^), somatic cells have successfully formed an epithelium with correct apical-basal polarity^19^ (Fig. S1A-B) while maintaining mitotic activity (Fig. 1H,I).

Another key difference in divisions of male CySCs and female FCs is their cytokinetic program. Male CySCs complete cytokinesis to release daughter cells from the niche^28^. By contrast, about half of female FC divisions occur with incomplete cytokinesis and formation of stable ring canals between daughter cells^27^. To address whether somatic loss of Chinmo induces the female-specific cytokinesis program, we first quantified cytokinesis in male CySCs. By visualizing CySCs from mitotic entry (as assessed by spindles labeling via tubulin::GFP) through abscission (as assessed by midbody labeling via anillin::RFP), we determined that male CySCs always complete cytokinesis within 3.5 hrs (Fig. 1J,M). This is consistent with measurements of abscission timing in other cell types^29,30^. Previous work and our own analyses find that incomplete cytokinesis in female FCs leads to retention of a midbody ring between daughter cells for longer than 3.5 hrs (Fig. 1K). Therefore, we quantified cytokinetic timing in 0-1D *chinmo*^*RNAi*^ testes and defined any somatic division retaining a midbody for *longer* than 3.5 hrs between daughter cells as an incomplete cytokinetic event. Interestingly, in 0-1D *chinmo*^*RNAi*^ testes, we find that a significant proportion of dividing somatic cells exhibit incomplete cytokinesis both within the niche (≤ 20.37 μm from the center of the niche) and outside the niche (> 20.37 μm from the center of niche) (Fig. 1L-N). Some of these *chinmo*-depleted FC-like cells retained midbodies at the intercellular bridge for 10-20 hrs (Fig. 1M). Importantly, 51% of ectopically dividing somatic cells exhibited incomplete cytokinesis in *chinmo*^*RNAi*^, which closely matches the proportion of FC divisions resulting in incomplete cytokinesis within epithelia of early egg chambers (57%)^27^. Thus, somatic loss of Chinmo induces a rapid conversion of testis somatic cells to a female-specific division and cytokinetic program that closely mimics the tight transition of female somatic cells from pre-follicle stages to bona fide follicular epithelium.

In non-epithelialized cells, constriction of the actomyosin contractile (AMC) ring is symmetric, resulting in central positioning of the midbody and central spindle^31^. This symmetric constriction is evident in male CySCs of the testis (Fig. S2A,E). By contrast, all epithelial cells asymmetrically constrict the AMC ring from the basal to apical surface, ensuring proper formation of apical adhesions between cells and retention of epithelial integrity ^29,32^. In female FCs of the ovary, this asymmetric constriction occurs with stable midbodies always localized to the apical surface of the epithelium^32^ and central spindle microtubules displaced from the center of the cell (Fig. S2B,E). Interestingly, somatic cells from 0-1D *chinmo*^*RNAi*^ testes exhibit a mixture of symmetric and asymmetric AMC ring constriction (Fig. S2C.E), demonstrating their progressive conversion to female-biased cellular behavior. Moreover, with 5 days of *chinmo* inhibition, all somatic cells residing in a complete epithelium exhibit asymmetric constriction of the AMC ring indistinguishable from female FCs (Fig. S2D,E), suggesting full conversion to female-specific cytokinetic programming. Apical localization of midbodies can be seen in cross sections of both female and *chinmo*^*RNAi*^ epithelia (Fig. S2F-G). Thus, by leveraging our powerful imaging system, we have identified the first appearance of female-specific behaviors during loss of somatic male sex identity.

### Sex-converted soma perform collective rotation similar to female follicle cells

A defining feature of sexually dimorphic gonads is how the somatic populations interact with the germline to promote differentiation. In testes, two somatic cells must fully encyst amplifying germ cells (Fig. 2A) as they co-differentiate to eventually produce sperm^11–15^. Encysting somatic cells move slowly (0.05 μm/min) down the coil of the testis (Fig. 2A’,G) following paths with no consistent directionality (Fig. 2D) and driven by their tight association with germ cells. In ovaries, FCs divide to form an epithelium around germ cell clusters and collectively migrate around the developing egg chamber (Fig. 2B) to promote oocyte elongation^16–18^. In our live imaging of early egg chambers, we find that FCs migrate in a cohesive, directed path at an average speed of 0.215 μm/min (Fig. 2B’,G,E), consistent with existing literature^16^. Given that *chinmo*-deficient somatic cells in the testis engage female-specific cytokinetic behaviors, we investigated whether the FC-like epithelium also initiates female-specific collective cell migration. Excitingly, our live imaging reveals that 5D *chinmo*^*RNAi*^ testis soma indeed engage functional rotation at a similar rate to female FCs (0.158 μm/min; Fig. 2C’,G). Importantly, this migration is collective and directional in the x-axis (Fig. 2F), a phenotype we never observe in somatic cells in wildtype testes.

**Figure 2.**
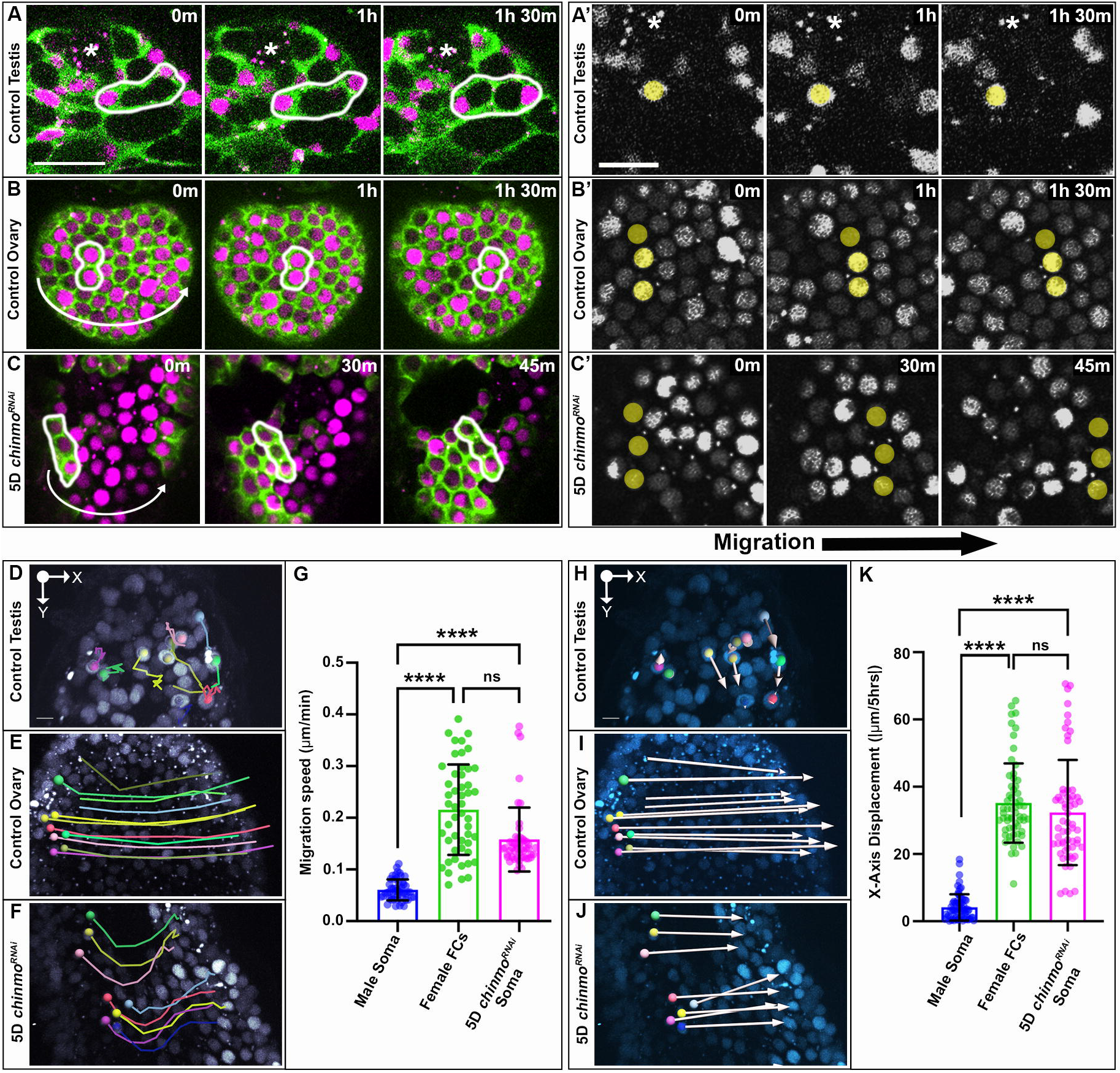
The sex-converted soma performs collective rotation similar to female follicle cells. (A-C) Time-lapse imaging of somatic expression of UAS tubulin::GFP (green) and UAS anillin::RFP (magenta) in a (A) control testis, (B) control ovary, and (C) 5D *chinmo*^*RNAi*^ testis. White outlines circle groups of cells that are either non-migratory or migratory. Scale bar: 20 μm. Curved arrows indicate directional migration of cells. (A’-C’) Time-lapse imaging of somatic nuclei (grey) in a (A’) control testis, (B’) control ovary, and (C’) 5D *chinmo*^*RNAi*^ testis. Yellow transparent dots indicate cell migration over time. Scale bar: 5 μm. Asterisks indicate stem cell niches. Each image is 1-2 z-slices. (D-F) Stills from live imaging displaying colored tracks that indicate the movement of nuclei over time in a (D) control testis, (E) control ovary, and a (F) 5D *chinmo*^*RNAi*^ testis. (K) Quantification of total x-axis displacement measured in |μm/5hrs| (n ≥ 60 cells in at least 4 samples). ****p<0.0001 (One-way ANOVA). ns, not significant. Error bars: standard deviation of the mean. All experiments n ≥ 2 trials. Scale bar: 5 μm (for D-F, H-J). Each Imaris image is 40 z-slices.

To further demonstrate the migratory behavior of *chinmo*^*RNAi*^ somatic cells, we used Imaris to generate total x-axis displacement over 5 hrs (Fig. 2H-J,K). By positioning an X/Y axis on the niche such that the y-axis runs along the posterior of the testis and x-axis runs perpendicular to the anterior-posterior axis, we calculated the total x-axis displacement of somatic cells. Unlike male somatic cells, which exhibit mostly y-axis directionality (Fig. 2H) and very small x-axis displacement (Fig. 2K), both female and *chinmo*-deficient somatic cells exhibit large x-axis displacement (Fig. 2I-K). Altogether, these findings represent the first evidence of sex-converted somatic cells performing functional female behaviors and broadens our understanding of how female-specific genetic programming can instruct behavioral changes.

### Somatic expression of adherens junction and ECM proteins is required for rotational migration

Adherens and extracellular matrix (ECM) proteins are known for facilitating apical-basal polarity in a variety of epithelial tissues^29^. For instance, the somatic epithelium in ovaries and *chinmo*-depleted testes express E-cadherin (Ecad) apically towards germ cells and secrete Perlecan (Pcan) basally towards the muscle sheath^33,34^. Prior studies have found that Pcan is essential for maintaining the integrity of the female follicular epithelium. Moreover, RNAi depletion of Pcan in a *chinmo*^*RNAi*^ background was shown to prevent full feminization of testes^34^. In fact, recent work has shown that disrupting the matrix protease AdamTS-A causes defects in the basement membrane deposited by FCs and severely disrupts rotational migration of the epithelium^35^. However, it is unknown how disrupting either adherens or ECM proteins within somatic cells can affect the ability of feminized testis soma to engage rotational migration.

The expansion of Fasciclin 3 (Fas3) expressing septate junctions (equivalent of tight junctions) between somatic cells has been used previously to quantify degree of feminization upon loss of Chinmo^19,21^. In wildtype testes, FasIII is enriched strictly at niche cell membranes and is absent from differentiating somatic cells (indicated by somatic-specific transcription factor Traffic jam (Tj); Fig. 3A). Somatic depletion of Chinmo from testes for 10-12D results in ectopic Fas3 expression in nearly all Tj-positive cells (Fig. 3B,D), demonstrating full acquisition of female-specific characteristics. Interestingly, somatically depleting Ecad in a *chinmo*^*RNAi*^ background prevents full feminization of testes (Fig. 3C-D), similar to decreased feminization observed upon combined loss of Chinmo and Pcan^34^. Together, these data suggest that adherens junctions and production of ECM components are critical for acquisition of female-specific somatic identity. Therefore, using live imaging and analyzing tracks (Fig. 3E-I), migration speed (Fig. 3K), and displacement (Fig. 3L-R), we evaluated whether Ecad and Pcan are required in FCs in the ovary and *chinmo*-deficient somatic cells in the testis for efficient epithelial rotation. Whereas in control ovaries, FCs migrate in the x direction in a continuous, organized path (Fig. 3E) at an average speed of 0.215 μm/min (Fig. 3K), FCs expressing either *Ecad*^*RNAi*^ and *Pcan*^*RNAi*^ migrate in a disorganized manner (Fig. 3F-G) at slower average speeds of 0.072 μm/min and 0.095 μm/min, respectively (Fig. 3K). Consequently, these cells exhibit significantly smaller x-axis displacement (Fig. 3M-N,R). Again, somatic cells from 5D *chinmo*^*RNAi*^ testes migrate continuously in a parallel path along the x direction (Fig. 3H) at an average speed of 0.158μm/min (Fig. 3K) similar to female FCs. Additional somatic depletion of either Ecad or Pcan in *chinmo*^*RNAi*^ backgrounds led to less organized somatic migratory paths (Fig. 3I-J) and reduced average speeds of 0.106μm/min and 0.075μm/min (Fig. 3K). As a result, these cells displayed significantly smaller x-axis displacement (Fig. 3P-Q), strongly demonstrating lack of coordinated migration in the absence of adherens and ECM proteins. Therefore, knockdown of either adherens or ECM proteins in female FCs and FC-like somatic cells results in lowered performance of rotational migration. Together, the data suggests that both adherens and ECM proteins are required for proper somatic collective cell migration both in wildtype ovaries and somatic sex-converted testes.

**Figure 3.**
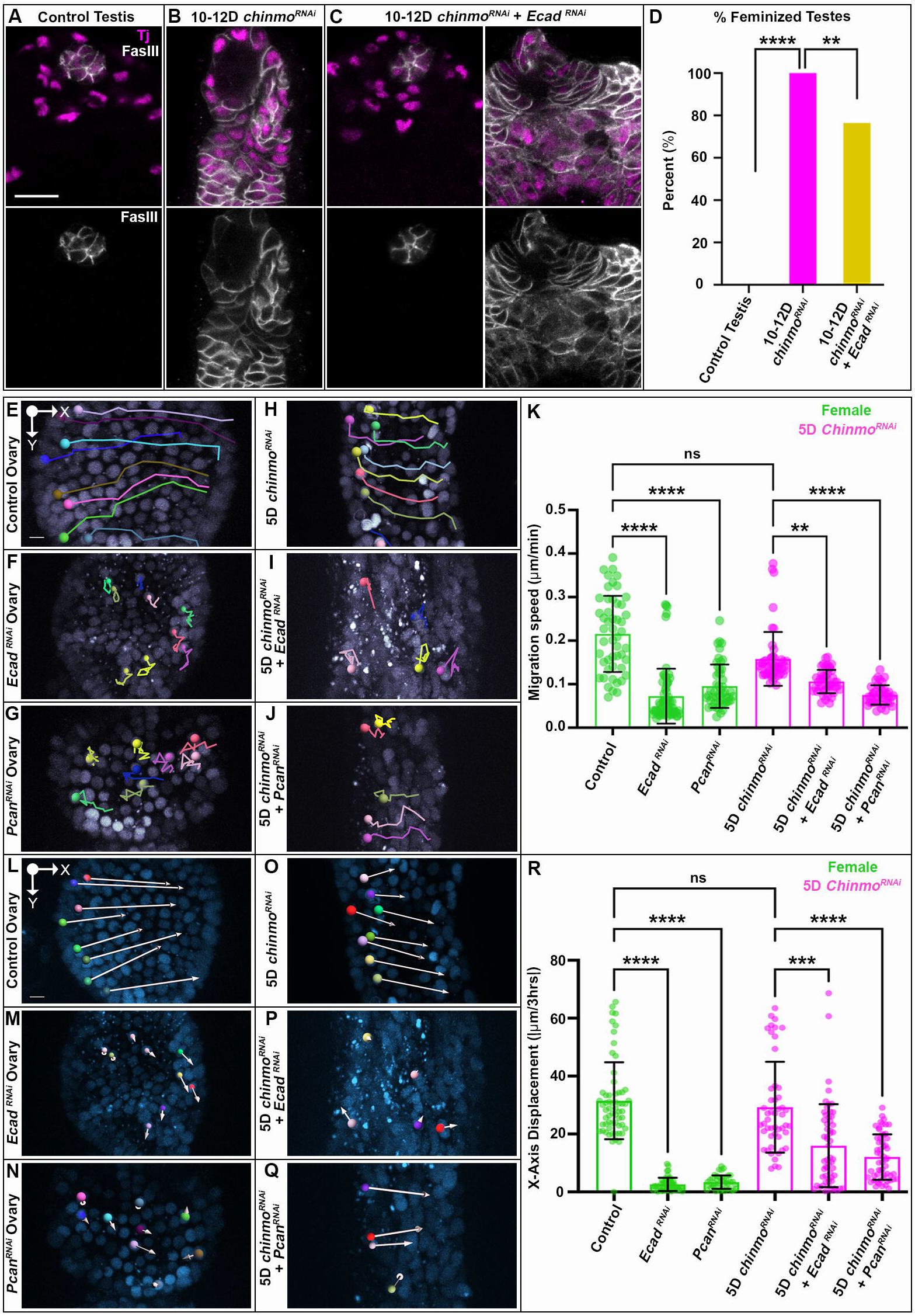
Somatic expression of adherens junction and ECM proteins is required for rotational migration. (A-C) Immunofluorescence staining of Traffic jam (magenta) and Fas3 (grey) in a (A) control testis, (B) 10-12D *chinmo*^*RNAi*^ testis, and in (C) 10-12D *chinmo*^*RNAi*^ + *Ecad*^*RNAi*^ testes. Scale bar: 20 μm. Each image is 1 z-slice. (D) Graph of percent feminized testes as measured by Fas3 expression in non-niche cells (*n* ≥ 15 testes). **p<0.0086, ****p<0.0001 (chi-squared test). (E-J) Stills from live imaging displaying colored tracks that indicate the movement of nuclei over time in a (E) control ovary, (F) *Ecad*^*RNAi*^ ovary, (G) *Pcan*^*RNAi*^ ovary, (H) 5D *chinmo*^*RNAi*^ testis, (I) 5D *chinmo*^*RNAi*^ + *Ecad*^*RNAi*^, and a (J) 5D *chinmo*^*RNAi*^ + *Pcan*^*RNAi*^ testis. (K) Quantification of migration speed in Quantification of migration speed in μm/min (*n* ≥ 40 cells in at least 8 samples). Data for control ovary and 5D *chinmo*^*RNAi*^ repeated from Fig. 2G. (L-Q) Live stills displaying x-axis displacement of somatic nuclei over 3 hrs in a (L) control ovary, (M) *Ecad*^*RNAi*^ ovary, (N) *Pcan*^*RNAi*^ ovary, (O) 5D *chinmo*^*RNAi*^ testis, (P) 5D *chinmo*^*RNAi*^ + *Ecad*^*RNAi*^, and a (Q) 5D *chinmo*^*RNAi*^ + *Pcan*^*RNAi*^ testis. (R) Quantification of total x-axis displacement measured in |μm/3hrs|. **p<0.0047, ***p<0.0006, ****p<0.0001 (One-way ANOVA). ns, not significant. Error bars: standard deviation of the mean. All experiments n ≥ 2 trials. Scale bar: 5 μm (for E-J, L-Q). Each Imaris image is 40 z-slices.

### Sex-converted soma induces early oocyte specification in XY germ cells

Although significant changes in the testis soma lacking Chinmo have been reported, accompanying differences in underlying XY germ cells remains to be explored. Rotation of the female follicular epithelium has been shown to cause a barrel-like rotation of germ cells within developing egg chambers^17,36^. By combining somatic and germline markers in a *chinmo*^*RNAi*^ background, we performed extended live-cell imaging to investigate changes in germ cell behaviors beneath the rotating feminized soma. Excitingly, we find that the *chinmo*-deficient, FC-like epithelium induces rotation of encapsulated germ cells (Fig. 4A-C), which closely mimics female egg chamber rotation. Interestingly, many *chinmo*-depleted testes exhibit partitioning of germ cells into clusters reminiscent of female egg chambers (Fig. S1B). Taken together, our data shows altered germ cell behaviors upon soma-specific manipulation of sex identity, begging the question of whether these changes in germline behavior also coincide with gain of female identity.

**Figure 4.**
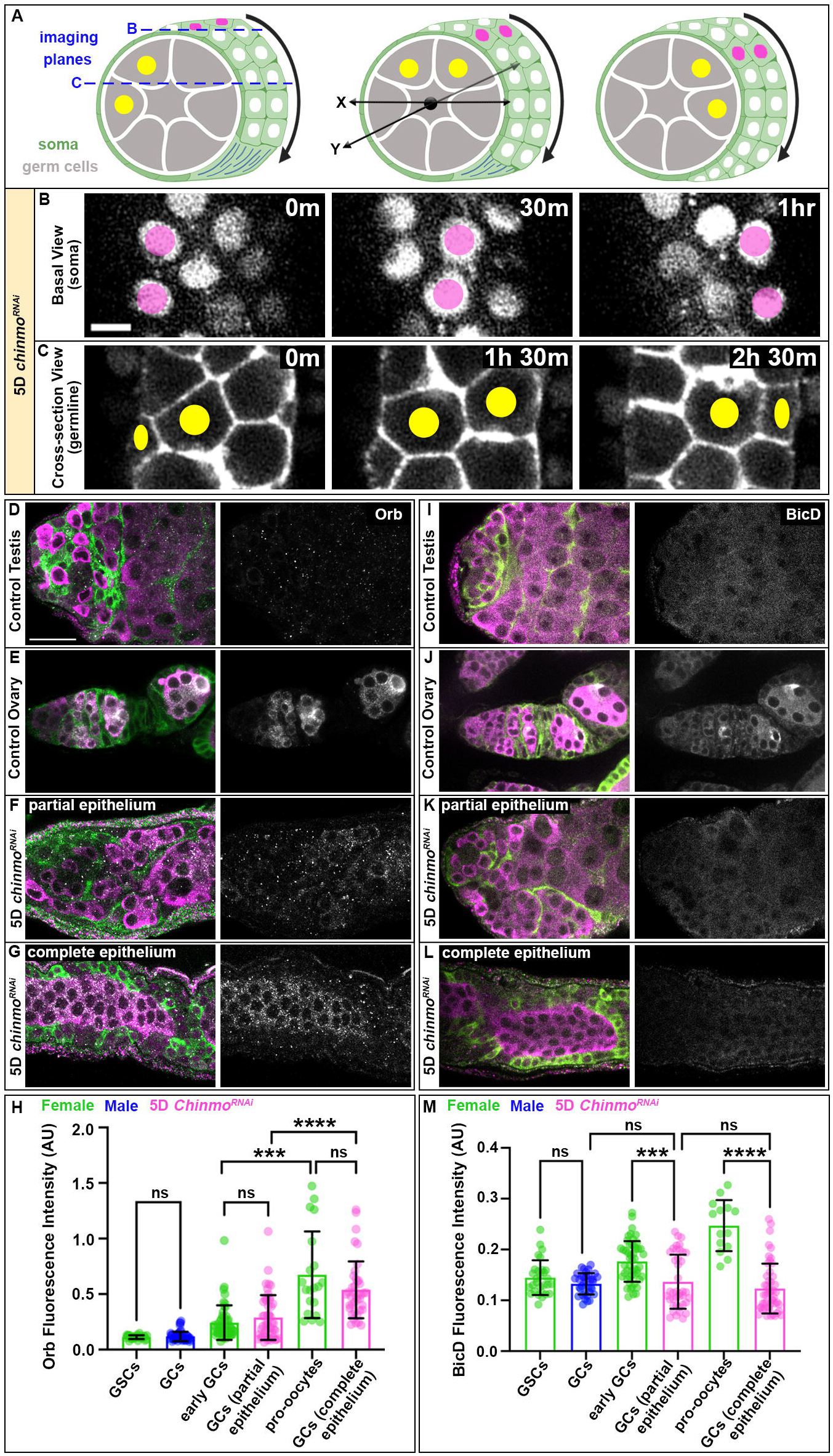
Sex-converted soma induce female-like behaviors and fate changes in the germline. (A) Diagrams of feminized somatic epithelium (magenta dots mark nuclei) rotating around germ cells (yellow dots mark nuclei). Blue dotted line indicates imaging planes. Black arrows indicate the direction of collective migration. Black x/y axis describes orientation of tissue. (B) Timelapse imaging of somatic anillin::RFP (grey) migrating in the x direction. (C) Timelapse imaging of germ cells expressing nos-lifeact::tdTomato (grey) migrating in the x direction. Scale bar: 5μm. (D-G) Immunofluorescent staining of somatic tubulin::GFP (green), germline Vasa (magenta), and Orb (grey) in a (D) control testis, (E) control ovary, (F) partial epithelium 5D *chinmo*^*RNAi*^ testis, and an (G) complete epithelium 5D *chinmo*^*RNAi*^ testis. (H) Quantification of Orb fluorescence intensity relative to Vasa (*n* ≥ 20 cells in at least 8 samples). (I-L) Immunofluorescent staining of somatic tubulin::GFP (green), germline Vasa (magenta), and BicD (grey) in a (I) control testis, (J) control ovary, (K) partial epithelium 5D *chinmo*^*RNAi*^ testis, and an (L) complete epithelium 5D *chinmo*^*RNAi*^ testis. (M) Quantification of BicD fluorescence intensity relative to Vasa (*n* ≥ 14 cells in at least 8 samples). ***p<0.0006, ****p<0.0001 (One-way ANOVA). ns, not significant. Error bars: standard deviation of the mean. All experiments *n* ≥ 2 trials. Scale bar: 20 μm (for D-G, I-L). Each image is 1-3 z-slices.

Recent work in the field has shown that the XY germline may be undergoing transcriptional changes when the somatic population is depleted of Chinmo^22^. This, along with our data, warrants investigation into whether these germ cells are specifying oocyte identity. One of the earliest markers of oocyte identity is the cytoplasmic polyadenylation element binding protein, Orb, which targets several mRNAs encoding proteins implicated in oocyte specification^37^. Immunostaining of Orb in control ovaries shows accumulation at low levels in the germline stem cells (GSCs), indistinguishable from levels observed in germ cells (GCs) from control testes (Fig. 4D-E,H). Consistent with prior literature, we show that in ovaries, Orb induction begins in early GCs and is significantly increased by the pro-oocyte stage (Fig. 4E,H). Importantly, the phenotype of 5D *chinmo*^*RNAi*^ is variable; while some testes exhibit a partial epithelium containing somatic cells still showing male morphology of long cytoplasmic extensions, other testes exhibit a cohesive or complete FC-like epithelium. Remarkably, Orb is induced in GCs from *chinmo*-depleted testes with a partial epithelium similar to early female GCs in the ovary (Fig. 4E-F,H) and is significantly increased to pro-oocyte levels in GCs enclosed by a complete epithelium (Fig. 4E,G-H). To our knowledge, this is the first indication that oocyte specification is, at least in part, non-autonomously controlled by the surrounding soma.

Furthermore, Orb staining always appears diffuse across all GCs in 5D *chinmo*^*RNAi*^ testes (Fig. 4G)—unlike in ovarian GCs, which quickly restrict Orb protein to the oocyte by the time the first egg chamber is formed. Bicaudal D (BicD) is another oocyte-specific protein that accumulates at the same time as Orb and is known for polarizing the oocyte microtubule cytoskeleton to restrict meiosis to the oocyte^38,39^. Immunostaining for BicD showed normal localization in ovaries (Fig. 4J), consistent with previous work^39^. However, we found no change in BicD levels in GCs from control testes and 5D *chinmo*^*RNAi*^ testes, suggesting no change in BicD expression induced by feminized somatic cells in the testis (Fig. 4I-M). Despite non-autonomous induction of Orb in the germline of somatically *chinmo*-depleted testes, the lack of enrichment of germ cell intrinsic proteins important for Orb restriction to the oocyte may explain why Orb levels appear to be equivalent across all GCs. In turn, this suggests that there are limits to progression to female identity in adult XY germ cells surrounded by *chinmo*-deficient somatic cells, which is consistent with prior work in developing gonads^40–43^.

Nevertheless, these findings uncover a novel role of the adult female soma in regulating female sex identity in adult germ cells. Finally, utilizing mismatched soma-germline sex identity in this model system, we are poised to begin testing soma-derived versus germline-intrinsic requirements for oocyte specification and identifying new molecular components of female-specific behaviors.

## Materials and Methods

### Experimental model and subject detail

*Drosophila melanogaster* stocks were maintained on Bloomington *Drosophila* Stock Center (BDSC) standard cornmeal medium in vials or bottles. All crosses were kept at 25°C unless otherwise indicated. Fly stocks included: Traffic Jam Gal4 (Kyoto Stock Center; nanos-lifeact::tdTomato^44^; UAS-chinmo-RNAi (BDSC #33638). UAS-Scra::mRFP (BDSC #52220); UAS-tubulin::GFP^11^; UAS-Ecad RNAi (BDSC #38207); UAS-Pcan RNAi (VDRC #24549).

### Time lapse imaging

Extended time-lapse imaging and culture conditions were adapted from those previously described^11,45–47^. Age matched samples were dissected in Ringers solution and mounted onto a poly-lysine-coated coverslip at the bottom of an imaging dish (MatTek). Ringers solution was removed and imaging media (15% fetal bovine serum, 0.5X penicillin/streptomycin, 0.2 mg/ml insulin in Schneider’s insect media) was added. Samples were imaged every 15 or 30 min for up to 24 hrs on an Olympus iX83 with a Yokagawa CSU-10 spinning disk scan head, 60X 1.4 NA silicon oil immersion objective and Hamamatsu EM-CCD camera using 1 μm z-step size (40 μm stacks). Experiments were repeated a minimum of two times and at least 7 samples were analyzed for each genotype/condition.

### Analysis of somatic cell behaviors from live imaging

To visualize somatic cell divisions, we somatically expressed UAS-anillin::RFP, which localizes to the nucleus and midbodies, and UAS-tubulin::GFP, which localizes to the cytoplasm and mitotic/central spindles. For quantification of the distance of dividing somatic cells, we calculated the distance between two points in 3D space (from the center of the niche to center of the dividing somatic cell). Therefore, the following formula was used: square root [(X mitotic cell – X niche)^2^ + (Y mitotic cell – Y niche)^2^ + (Z mitotic cell – Z niche)^2^]. A sample of at least 20 cells were tracked over time per sample.

To quantify somatic cell cytokinesis, we first identified male CySCs (indicated by close proximity of nuclei to the niche and cytoplasmic contact with the niche). Mitosis was determined by the presence of mitotic spindles. Cytokinesis progression was observed by central spindle formation and midbody condensation. Final abscission events were marked when the condensed midbody was significantly displaced from the intercellular bridge and engulfed by adjacent somatic cells as well as movement of daughter cell nuclei from one another. Retention of the midbody at the intercellular bridge for longer than 3.5 hrs was considered an incomplete cytokinesis event.

To determine symmetric versus asymmetric constriction of the somatic AMC rings, we first identified somatic cells undergoing constriction of the mitotic furrow. Using anillin to measure the furrow length and central spindle to measure the length from one end of the furrow to the central spindle, we first divided the length from one end of the furrow to the central spindle over the total length of the furrow. One half was subtracted from these values to represent displacement from the center of the furrow.

For quantification of somatic cell migration speed, the ImageJ manual tracking tool was used to track somatic nuclei over three consecutive time points. The parameters were set to 30 minute intervals with an x/y calibration of 4.6154mm and z calibration of 1. Values generated by this tool were then divided by 30 to get the distance (mm) traveled per minute. Visual displays of tracks were generated in Imaris 10.2 (Bitplane, Oxford, UK) using the spot function tool.

Imaris 10.2 spot algorithm was used to manually track somatic cells over 3- and 5-hr time lapses. Using a reference frame centered on the testis niche with the Y axis pointed posteriorly, total track displacement along the X axis reference frame was determined. If necessary, drift correction was implemented.

### Immunostaining

Immunostaining was performed as previously described^11,45,48^. In short, samples were dissected in Ringers solution and fixed for 30 min in 4% formaldehyde in Buffer B (75 mM KCl; 25 mM NaCl; 3.3 mM MgCl2; 16.7 mM KPO4) followed by multiple washes in PBSTx (1× PBS, 0.1% Triton-X 100) and blocking in 2% normal donkey serum. Samples were incubated in primary antibodies at 4°C at least overnight, washed multiple times, and then incubated in appropriate secondary antibodies for 1 hr at room temperature. After additional washes, samples were equilibrated in a solution of 50% glycerol and then mounted on slides in a solution of 80% glycerol. Primary antibodies used were: rat anti-DE-cadherin [Developmental Studies Hybridoma Bank (DSHB), 1:20], mouse anti-Orb (DSHB, 1:30), chicken anti-GFP (Aves Labs, 1020, 1:1000), guinea pig anti-Traffic jam (Dorothea Godt, University of Toronto, Canada, 1:5000), mouse anti-Fasciclin 3 (DSHB, 1:50), rabbit anti-Vasa (Boster Biological Technology Co, DZ41154, 1:5000), and mouse anti-BicD (DSHB, 1:100).

Secondary antibodies used were from Jackson ImmunoResearch and used at 1:125 dilution: Alexa fluor-488 (anti-chicken 703-545-155, anti-rat 715-545-151), -Cy3 (anti-guinea pig 706-165-153, anti-mouse 715-165-153, anti-rabbit 711-165-152) and -Cy5 (anti-rat 712-605-153, anti-mouse 715-605-151). All antibodies have been previously verified by the Drosophila community.

### Quantification of fluorescence intensities

For analysis of Orb and BicD induction, mean fluorescence intensities were quantified within a single z slice through the center of germ cells and ratioed to the mean fluorescence intensities of Vasa. For female samples, three GSCs, 5 early germ cells, and 2 pro-oocytes were measured per ovary. For male samples, 5 germ cells were measured per testis. Thus, the following formula was used: (mean Orb/BicD – mean background Orb/BicD)/(mean Vasa – mean background Vasa).

All images of fixed and immunostained testes were acquired using a Leica Stellaris 5 DMi8 inverted stand with tandem scanner; four power HyD spectral detectors; and HC PL APO 63×/1.4NA CS2 oil objective using LAS X software.

### Analysis of Fas3

Testes were raised at 18°C until eclosion and then upshifted to 29°C for maximal Tj-Gal4 activity. After 10-12 days at 29°C, testes were dissected and stained for Fas3, Tj and Vasa as described above, and then mounted in Vectashield with DAPI (Vector laboratories, Catalog # H-1200). Testes were scanned on a Zeiss 700 confocal microscope at 63x. Testes were then monitored for the expression of Fas3 in non-niche cells. As a reference, 0% of control testes and 100% of testes somatically depleted for Chinmo display Fas3 outside of the niche. Somatic co-depletion of Chinmo and ECad resulted in a significant decline in the percentage of testes displaying Fas3 in non-niche cells as assessed by chi-square test.

### Quantification, statistical analysis and image processing

Time-lapse images were analyzed and z-projections generated using ImageJ software. All graphical representations of data and statistical analysis were performed in Graphpad Prism (One-way ANOVA and non-parametric Mann–Whitney U-test). Error bars represent standard deviation. n and P values are indicated in figure legends. Fixed, end-point analyses were based on previous analyses in the field (Fairchild et al., 2016; Li et al., 2014). Figures were generated using BioRender.com and Adobe Photoshop.

