## Supplemental Figures for "Conversion of somatic sex identity in the testis induces female-specific cellular behaviors in the soma and early oocyte specification in the germline"

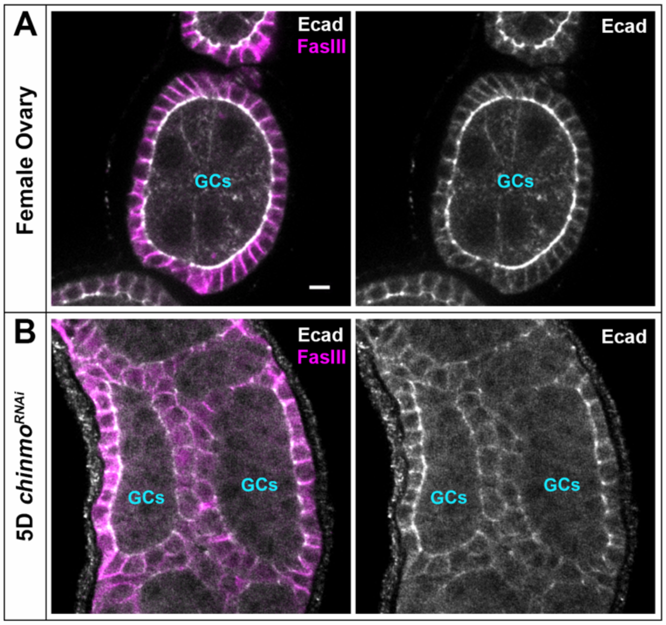


**Supplemental Figure 1.**
(A-B) Immunofluorescent staining of Ecad (grey) and Fas3 (magenta) in a (A) control ovary and (B) 5D *chinmo*$^{RNAi}$ testis, both showing apical localization of Ecad towards germ cells. Germ cells (GCs) are indicated. Scale bar: 5 µm. Each image is 1 z-slice.

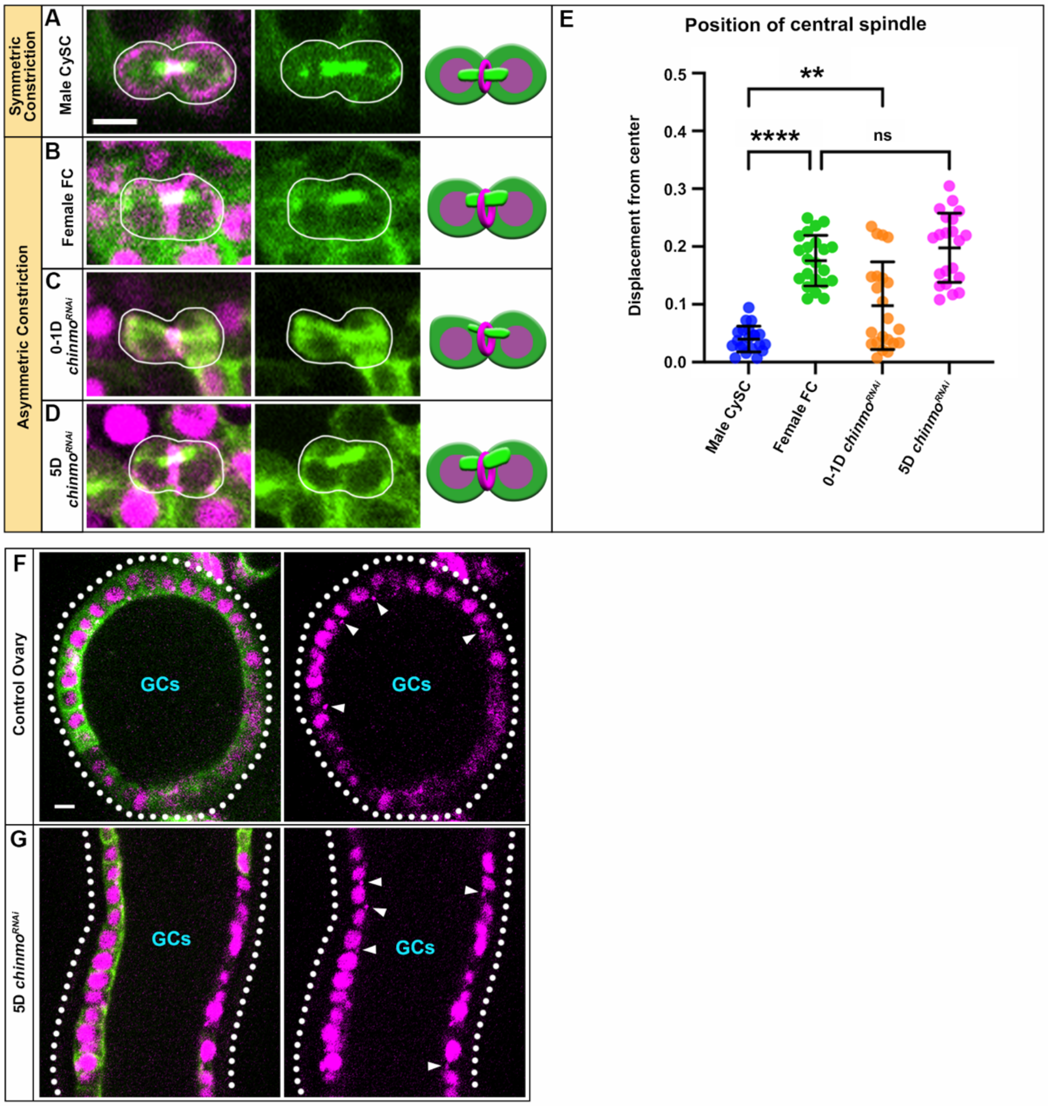


**Supplemental Figure 2.**
(A-D) Stills from live imaging of somatic tubulin (green) and anillin (magenta) during constriction of the AMC ring in a (A) male CySC, (B) female FC, (C) 0-1D *chinmo*$^{RNAi}$ somatic cell, and (D) 5D *chinmo*$^{RNAi}$ somatic cell. (E) Quantification of displacement of the center of the furrow ($n \geq 21$ cells in 7 samples). **p<0.0047, ****p<0.0001 (One-way ANOVA). Ns, not significant. Error bars: standard deviation of the mean. (F-G) Stills from live imaging of somatic tubulin (green) and anillin (magenta) showing apical localization of midbodies (arrowheads) in a (F) control ovary and (G) 5D *chinmo*$^{RNAi}$ testis. Dotted outline marks basal periphery of the tissue. Germ cells (GCs) are indicated. All experiments $n \geq 2$ trials. Scale bar: 5µm (for A-D, F-G). Each image is 1-4 z-slices.
